# The first fossil replete ant worker establishes living food storage in the Eocene

**DOI:** 10.1101/2022.12.15.520604

**Authors:** Indira Sawh, Eunice Bae, Luciana Camilo, Michele Lanan, Andrea Lucky, Henrique Morais Menezes, Gianpiero Fiorentino, Christine Sosiak, Lily Khadempour, Phillip Barden

**Affiliations:** Department of Earth and Environmental Sciences, Rutgers University, Newark, New Jersey, USA; Department of Biological Sciences, New Jersey Institute of Technology, Newark, New Jersey, USA; Graduation Program in Natural Sciences/Biology, Federal University of Maranhão, Imperatriz, Brazil; Department of Entomology, University of Arizona, Tucson, Arizona, USA; Entomology & Nematology Department, University of Florida, Gainesville, Florida, USA; Postgraduate Program in Environmental Sciences (PPGCAM), Federal University of Maranhão, Chapadinha, Brazil; Division of Invertebrate Zoology, American Museum of Natural History, New York, New York, USA

**Keywords:** Palaeoentomology, repletism, *Leptomyrmex*

## Abstract

Worker specialization extends the behavioral and ecological repertoire of ant colonies. Specialization may relate to colony defense, brood care, foraging, and, in some taxa, storage. Replete workers swell the crop and gaster to store liquid food, which can be accessed by other colony members through trophallaxis. This storage ability, known as repletism, has independently evolved across several ant lineages, but the temporal history of this trait has not yet been investigated. Here, we describe the first fossil replete in the extinct species *Leptomyrmex neotropicus* Baroni Urbani, 1980 preserved in Miocene-age Dominican amber. Together with new evidence of repletism in *L. neotropicus’* extant sister species, *L. relictus* Boudinot et al., 2016, we reconstruct the pattern of acquisition and descent in this storage-linked trait. Our ancestral state reconstruction suggests that *Leptomyrmex* acquired replete workers in the Eocene and may therefore represent the earliest instance of so-called “honeypot” ants among all known ants, both living and extinct.

## Introduction

Eusociality is a profound phenotypic phenomenon that shapes morphology as well as behavior. Division of labor is central to advanced sociality in insects and workers may exhibit a range of behavioral or morphological specializations related to task performance (Wilson & Hölldobler 2005). A striking example of caste specialization is repletism. Replete workers serve as living food storage within the colony by retaining liquid food within their gaster (terminal abdominal segments). Food storage takes place in the crop, a region of the alimentary canal in the foregut between the esophagus and the proventriculus. The crop swells to accommodate large amounts of food, distending the gaster to large proportions via elastic intersegmental membranes located between each tergite (Carney 1969, Wilson 1974). This elasticity enables repletes to distend their gaster dramatically in some species, which renders replete individuals visibly distinct from other workers (Conway 1994) and may limit mobility (Charbonneau & al. 2017). During times of scarcity, the stored contents of the replete crop are redistributed to colony members. Food is regurgitated from one ant to another, a process known as trophallaxis. Prior to trophallaxis, ants concentrate the stored material by reducing its water content and they add components of their internal fluids in the crop (Meurville & LeBoeuf 2021). This creates a network of fluid and nutrient exchange in the colony. Because trophallaxis is a common feature across ant lineages, many taxa have the capacity to distend the crop and gaster as part of a colony-wide “social stomach” (Meurville & LeBoeuf 2021). Taxa with a replete caste are ostensibly less vulnerable to fluctuating resource availability, particularly during seasons when food sources are limited (Van Elst & al. 2021). Trophallaxis may regulate the flow of nutrients among the colony with repletes, in particular, providing a reliable source of a “higher quality” of food (Borgesen 2000).

Repletism is a convergently evolved trait that has been observed in several ant lineages. While a precise definition of repletism is lacking, well-documented replete castes are reported from 20 genera (Andersen 2002, Borgesen 2000, Casadei-Ferreira & al. 2020, Cosens & Toussaint 1985, Glancey & al. 1973, Eyer & al. 2012, Conway 1992, Khalife & Peeters 2020, Lorinczi 2016, Moffett 1986, 1988, Ruano & Tinaut 1999). Repletes are typically classified as either crop repletes, storing liquid carbohydrates in the social stomach (i.e., crop), or fat body repletes, containing lipids in hypertrophied fat bodies (Charbonneau & al. 2017, Tschinkel 1987). Both types of repletes play a role in regulating nutrient storage in the colony and can provide sustenance to the colony during times of scarcity (Borgesen 2000, Charbonneau & al. 2017, Khalife & Peeters 2020). Crop repletes are often referred to as honeypot ants or honey ants, while fat body repletes are referred to as corpulents or “false honeypot ants” (Borgesen 2000, Lorinczi 2016). Nearly all origins of repletism occur in the subfamilies Formicinae and Myrmicinae, while a single dolichoderine genus – *Leptomyrmex* Mayr, 1862 – exhibits honeypot workers (Fig. 1).

**Figure 1.**
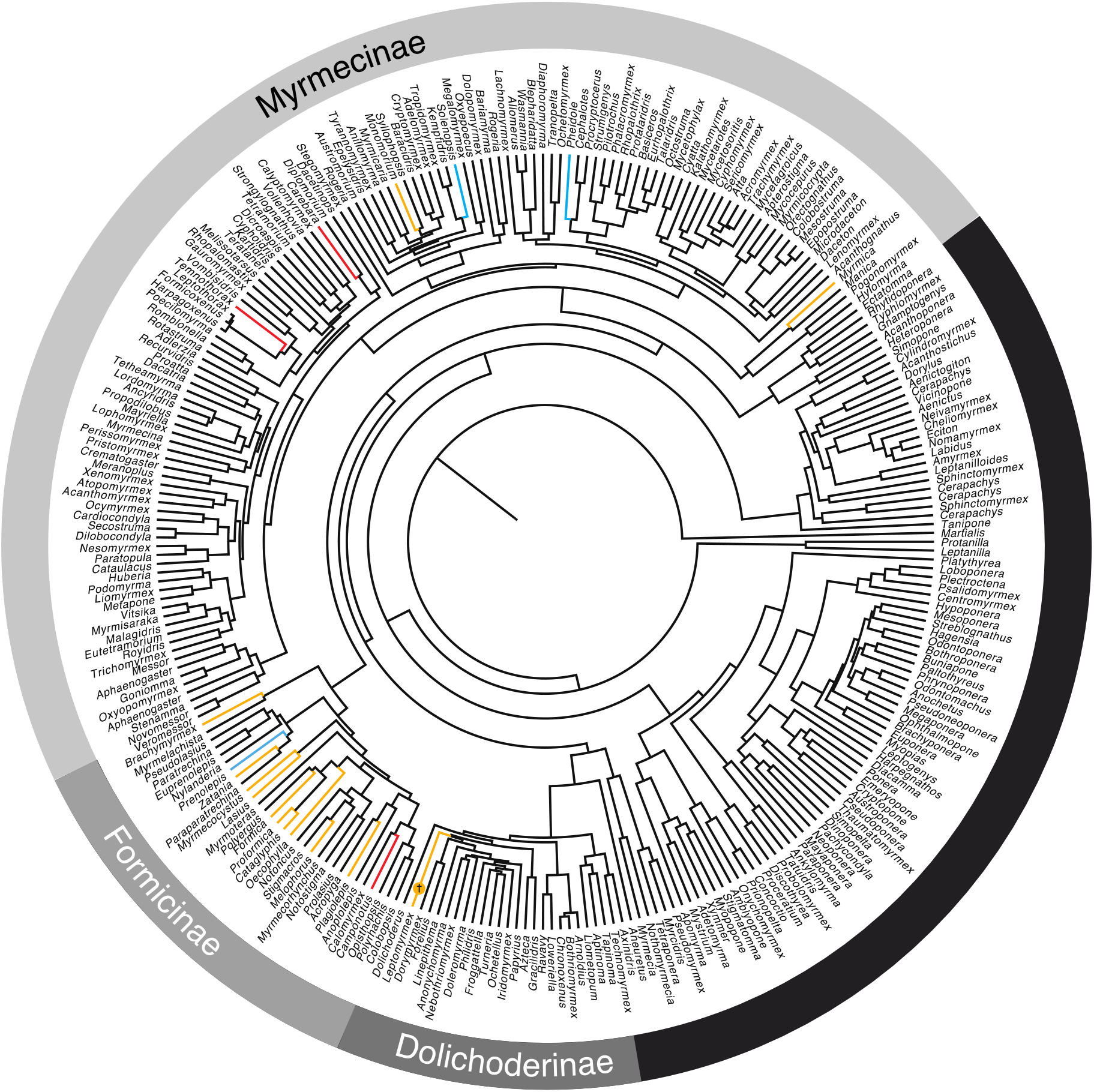
Phylogenetic distribution of known repletes. Topology adapted from Moreau and Bell, 2017. Lineage color reflects presence of repletes. Yellow: crop repletes; blue: fat body repletes; red: crop/fat body repletes. The fossil species *Leptomyrmex neotropicus* is denoted by the dagger † symbol.

While most of the 29 described *Leptomyrmex* species are endemic to Australia, New Guinea, and New Caledonia, two species are known from the neotropics: a single fossil species from the Dominican Republic and a recently discovered extant species in Brazil (Barden & al. 2017, Boudinot & al. 2016, Lucky & Ward 2010). Numerous Australasian *Leptomyrmex* species exhibit replete workers, which are frequently found outside of the nest and apparently use their distended crops for liquid food transport as well as storage (Wheeler 1915). Until now, the replete status of neotropical *Leptomyrmex* species has remained unknown, obscuring the temporal and biogeographic origin of this trait. Following the recent discovery of *L. relictus* in the Brazilian cerrado (Boudinot & al. 2016), *Leptomyrmex* is hypothesized to have originated in the Neotropics during the Eocene before dispersing to Australasia prior to the glaciation of Antarctica (Barden & al. 2017), a route that has been documented in other lineages (Dlussky & Radchenko 2013, Sanmartiin & Ronquist 2004). Were replete workers gained recently in Australasia or did repletes evolve in the Neotropics prior to their long-distance migration and diversification? Are there multiple origins of repletism in *Leptomyrmex?*

Here, we report new fossil and extant evidence of repletism in the Neotropics. Through microCT imaging we confirm the replete status of the now extinct Caribbean species *L. neotropicus* Baroni Urbani, 1980 and report replete workers in the extant sister species *L. relictus* Boudinot et al., 2016 for the first time. With these natural history data, we estimate the approximate age and retention of repletism in the genus *Leptomyrmex* through ancestral state reconstruction. Our approach illuminates the evolutionary history of extreme morphology-assisted food storage in ants.

## Materials and methods

### Fossil imaging

Photomicrographs were taken using a Nikon SMZ25 stereomicroscope equipped with a DS-Ri2 digital camera. Individual images were digitally stacked using Nikon NIS Elements to generate a high-resolution extended focus montage image. X-ray computed tomography data were generated at the New Jersey Institute of Technology Otto H. York Center for Environmental Engineering and Science using a Bruker SkyScan 1275 micro-CT scanner. The specimen was scanned at a voltage of 38kV and current of 190μA for 65ms exposure times averaged over four frames per rotation with a voxel size of 8.00μm. Z-stacks were generated using NRecon (Micro Photonics, Allentown, PA), segmented using 3D Slicer v4.9 (Fedorov & al. 2012), and rendered in Blender v.3.2.1.

### Ancestral state reconstruction

We reconstructed ancestral states of repletism across *Leptomyrmex* workers using the phylogeny of Barden & al. (2017) and a survey of natural history observations. Replete codings were derived from a literature survey as well as published iNaturalist accounts of reliably identified *Leptomyrmex* species (Supplemental Table 1). Given the uncertainty associated with some species regarding the presence of repletes, species were coded in a probability matrix: terminals were assigned a 1/0 replete/non-replete status if known to have repletes, a 0/1 replete/non-replete status if known to not have repletes, and assigned 0.5/0.5 if the presence of repletes was uncertain. We assumed a flat uninformative prior probability distribution for uncertain states rather than attempting to assess the probability of repletes vs non-repletes in uncertain species, given the lack of natural history information for many species. We conducted ancestral state reconstruction using stochastic character state mapping implemented with the prior probability matrix for character states. We inferred the reconstruction under the equal rates (ER) model, based on prior comparisons of the Akaike information criterion (AIC) using different character state evolution models (symmetrical (SYM) and all rates different (ARD), and ran the simulation for 200 trees (nsim=200)). State changes were summarized across all 200 trees. Ancestral state construction was conducted in R version 4.2.0 using the package phytools (Revell 2012).

## Results

### A fossil replete

Specimen BALDR-0155 is a *Leptomyrmex neotropicus* worker preserved as an inclusion within amber dated to the Upper Miocene (~16 Ma; Iturralde-Vincent & MacPhee 1996) from the Northern mines of near La Cumbre, Dominican Republic. The gastral elastic intersegmental membrane is significantly distended (Fig. 2) while there are no signs of taphonomic distortion across the cuticle. X-ray computed tomography recovers a sharp difference in inclusion density in the region of the gaster and head (Fig. 3), consistent with air. This heterogeneous density is the result of void space within the cuticle, a common feature recovered through X-ray imaging as internal features degrade after an insect is entombed in resin (Dierick & al. 2007).

**Figure 2.**
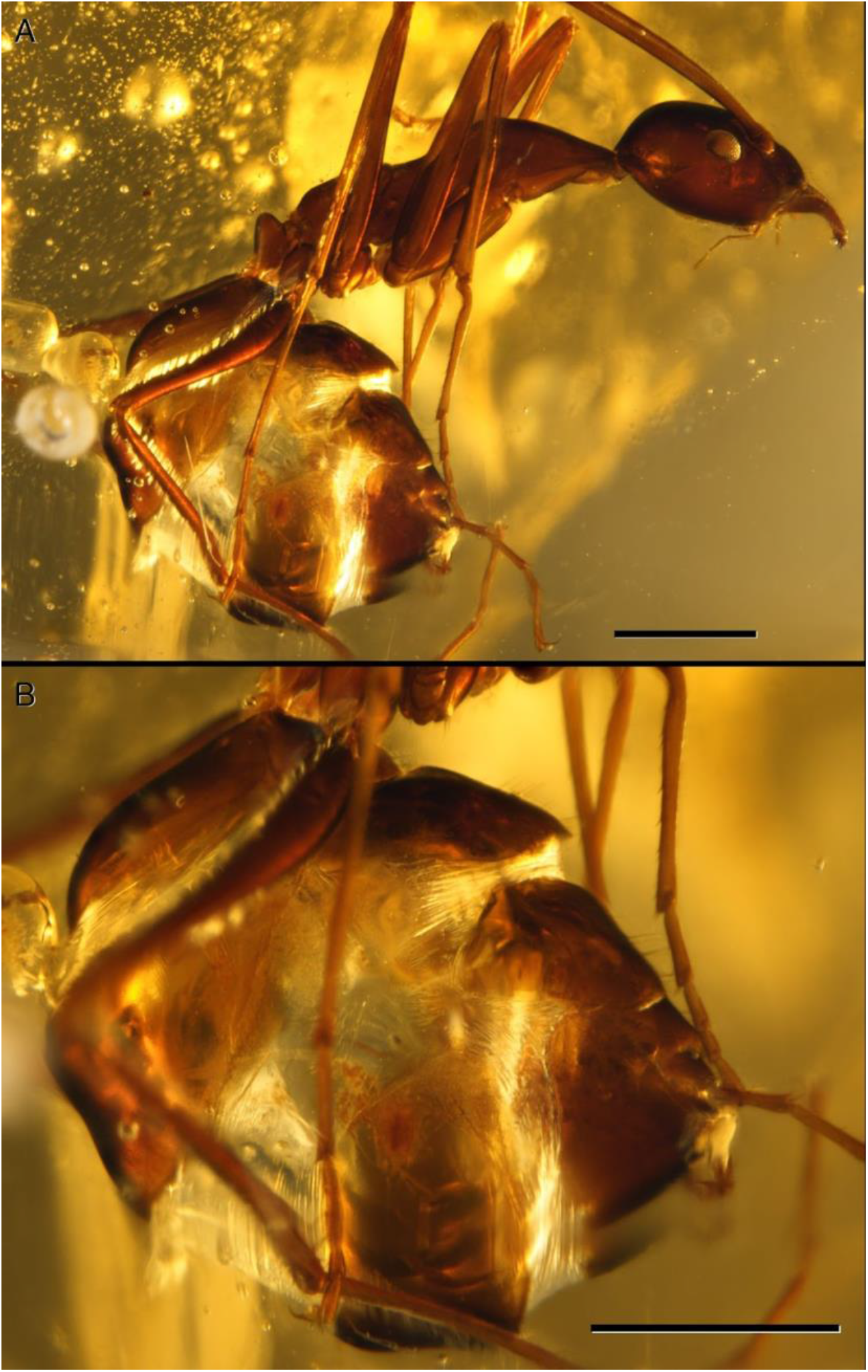
Photomicrograph of *L. neotropicus* replete specimen BALDR-0155 preserved in Miocene-age Dominican amber. (A) Lateral view. (B) Enlarged, lateral view of distended gaster. Scale = 1mm.

**Figure 3.**
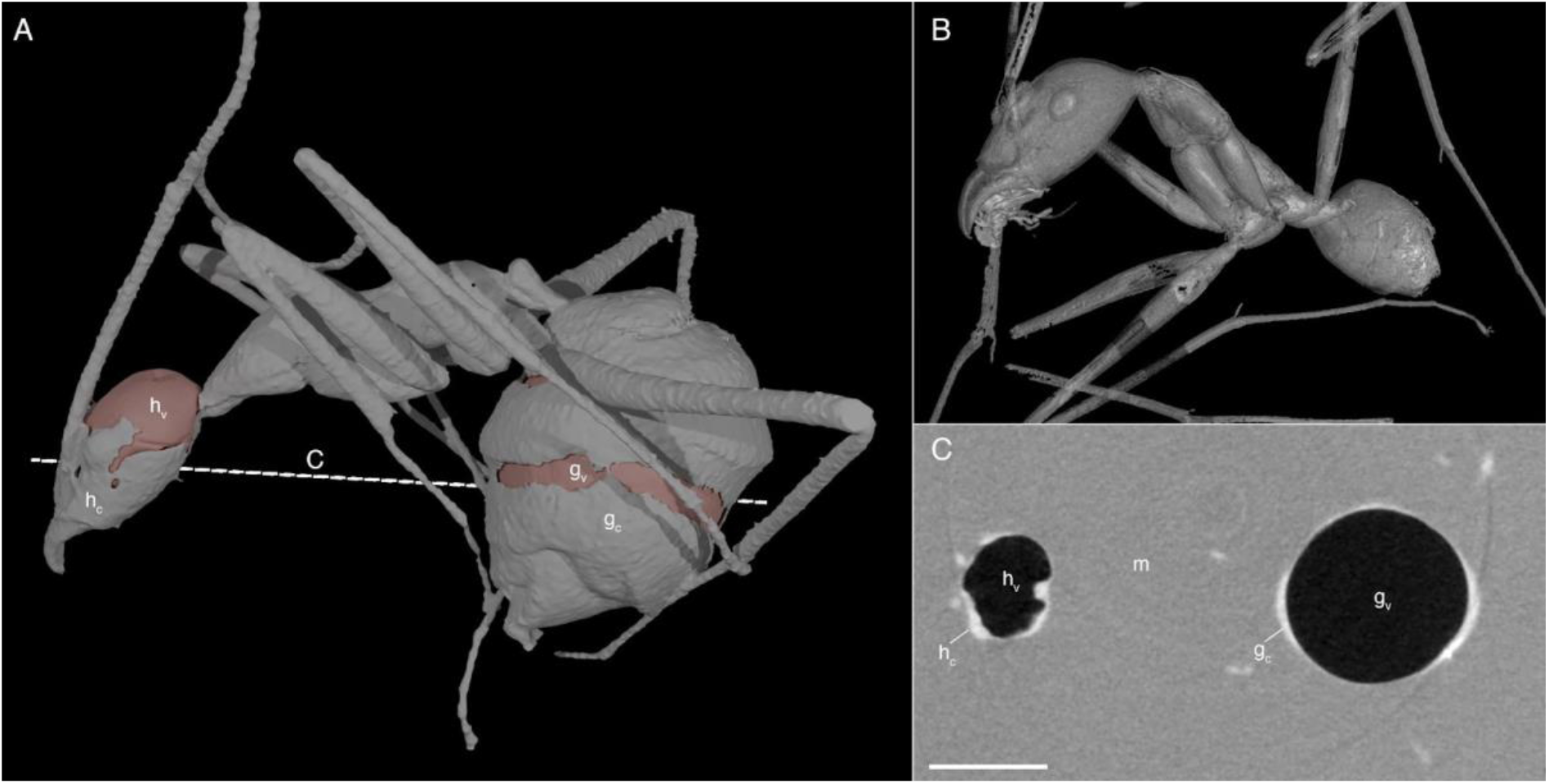
X-ray computed tomography images of *L. neotropicus*. (A) Lateral view of replete *L. neotropicus* worker specimen BALDR 0155. (B) Lateral view of a non-replete *L. neotropicus* worker (AMNHDR-13-85 modified from Barden et al., 2017). (C) Z-stack cross section of specimen BALDR 0155 head and gaster denoted by dotted line in sub-panel A. hc = head cuticle; hv = voidspace of head; m = amber matrix; gc = gaster cuticle; gv = voidspace of gaster. Scale = 1mm

### Extant repletes in the Neotropics

We (LC, HMM) observed replete workers of *Leptomyrmex relictus* entering and exiting a disturbed nest entrance, with some repletes carrying brood or unidentified objects in their mandibles (Supplemental Video 1 and 2). Replete gasters are conspicuously enlarged and distended relative to nearby non-repletes. This documentation in *L. relictus* confirms mobility and multiple task performance of repletes, as described in other *Leptomyrmex* species (Plowman 1981). Observations took place across the months of July, August, and September 2020 in Parque Cesamar (−10.209838, −48.322934) city of Palmas, state of Tocantins, Brazil. Two videos were recorded from the same nest within the park (Supplemental Video 1, 2).

### The evolution of repletism in *Leptomyrmex*

We found strong support for repletism as the ancestral condition of *Leptomyrmex* (Fig. 4; posterior probability 0.89 repletism). State changes were relatively infrequent; across all trees, we estimated the average number of gains and losses as 2.5. Our reconstruction suggests that once repletism evolved in *Leptomyrmex,* it infrequently or perhaps never reverted. The preponderance of ancestral nodes estimated as replete suggests that many *Leptomyrmex* species are likely to have a replete caste upon further study, though because our probability matrix used a flat uninformative prior due to lack of ecological data, this may have biased some more recent ancestral nodes towards repletism.

**Figure 4.**
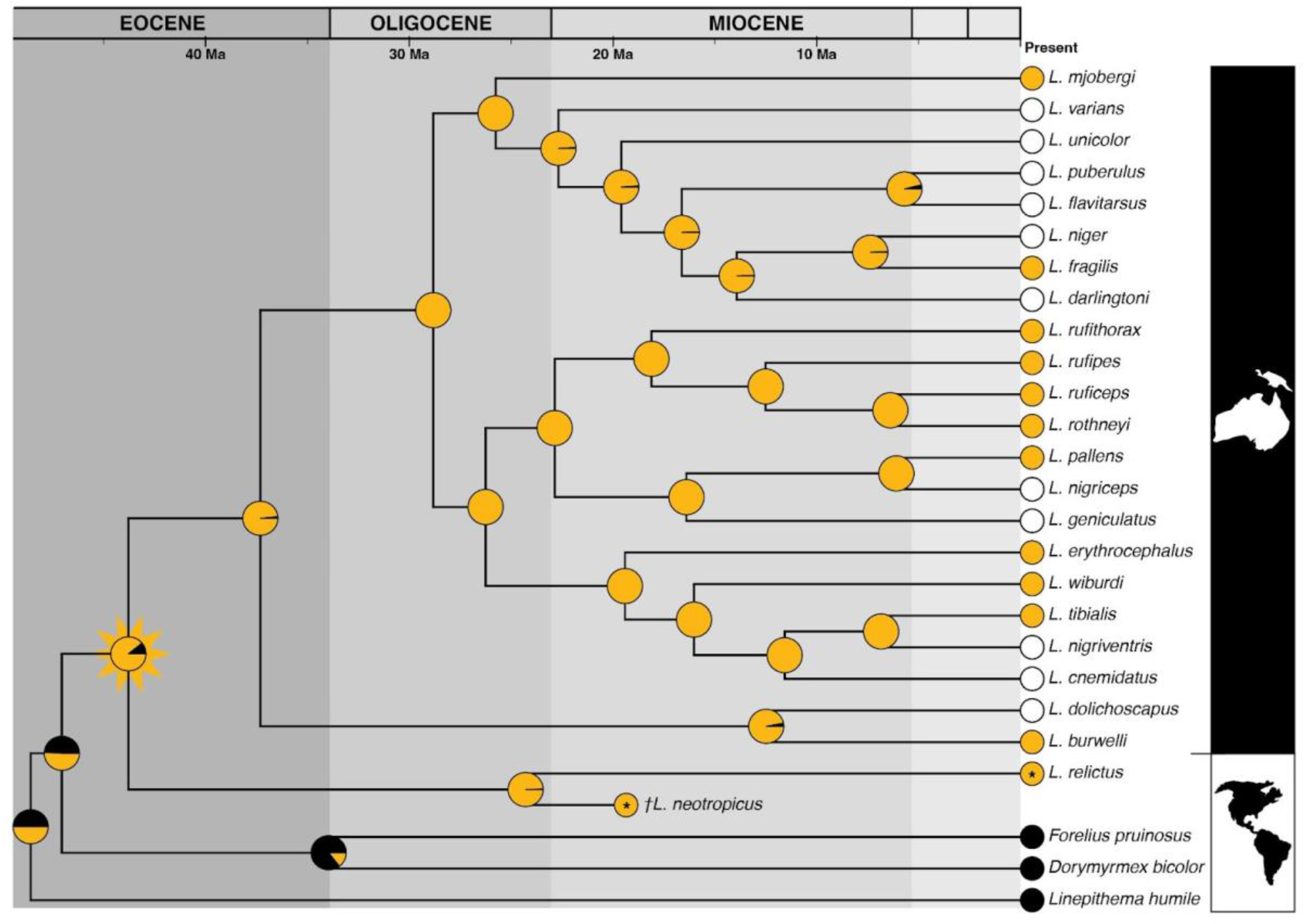
Ancestral state reconstruction of replete workers across *Leptomyrmex.* Summary of 200 simulated stochastic character histories under an equal rates (ER) model. Node pie charts represent posterior probabilities of states at each node. Yellow = replete; black = no repletes; white = unknown, these were coded as ambiguous in ancestral state reconstruction. Most recent common ancestor of all *Leptomyrmex* is indicated with star icon. Topology and mean node ages from Barden et al. (2017).

In extreme cases of repletism (e.g., *Myrmecocystus* Wesmael, 1838), replete workers tend to be immobile and confined to the nest, solely serving as subterranean food storage (Conway 1977). In fat repletes, once workers have depleted the resources in their fat bodies, usually during the season after storage, they also become foragers (Williams and Lucky 2020). In other taxa, repletes are mobile, performing other tasks, such as carrying brood or foraging (Cosens & Toussaint 1985, Conway 1992, Plowman 1981, Skinner 1980). Several species of *Leptomyrmex* are documented as mobile repletes, foraging on plants and transporting liquid food to the nest (Plowman 1981, Davidson & al. 2004). Our report of mobile repletes in *L. relictus,* and the presence of a *L. neotropicus* replete worker in fossil amber suggests that mobility and replete foraging was ancestral in this lineage.

## Discussion

We recover a single origin of replete workers in the last common ancestor of all extant and extinct *Leptomyrmex* species in the Eocene ~45 Ma (Fig. 4). Our results suggest that living food storage was present in a Neotropical ancestor and that this trait was retained as the genus expanded into Australasia. The expansion of grasslands and increases in global temperatures during the Eocene-Miocene transition may have contributed to the retention of repletism even across continents and tens of millions of years (Azevedo & al. 2020, Dlussky & Radchenko 2013). The retention of this trait in *Leptomyrmex* is unexpected because repletism is frequently ascribed to species that inhabit dry climates or are winter active (Hölldobler & Wilson 1990, Kronauer & al. 2004), while some replete species in *Leptomyrmex* are found in wet forests. Several other genera, including *Pheidole* Westwood, 1839 (Tsuji 1990), exhibit repletes within species that are endemic to wet habitats; even as climate is strongly linked to living food storage in some lineages, it does not appear to be a requirement for repletism.

It is notable that in our ancestral state reconstruction (Figure 4), and throughout ant lineages that contain repletism (Table 1), there remain many species with an unknown status. Repletism is often difficult to demonstrate if the replete workers are immobile and confined to the nest. These ants can be difficult to extract from underground, and nestmates often move repletes to deeper chambers to avoid exposure. There is therefore a bias toward underreporting replete castes, where they may exist, and it is more likely that fossilized lineages will exhibit evidence of mobile repletes since taxa with immobile repletes are unlikely to be aboveground and therefore caught in resin- or sediment-based preservation modes.

**Table 1.** Summary of major replete lineages and their estimated crown ages.

| Replete genera | Type of repletism | Mean lineage age (Ma) | Age reference | Replete reference |
| --- | --- | --- | --- | --- |
| *Agraulomyrmex* | Fat body | ~23 | Blaimer et al. (2016) | Prins (1983) |
| *Brachymyrmex* | Crop | ~17 [27.5 – 7.5] | Boudinot et al. (2022) | Zolessi et al. (1978) |
| *Camponotus* | Crop | ~24 [30 – 13] | Boudinot et al. (2022) | Lubbock (1880); Froggatt (1896) Heterick (2022) |
| *Cataglyphis* | Crop | ~15 (divergence) | Tinaut and Ruano (2021) | Eyer et al. (2013) |
| *Colobopsis* | Crop | ~24.5 [35 – 15] | Boudinot et al. (2022) | Wilson (1974); Hasegawa (1993) |
| *Lasius* | Crop | ~21.9 [28.6 – 15.3] | Boudinot et al. (2022) | Cammaerts (1996) |
| *Melophorus* | Crop | ~44.2 [51 – 20] (divergence) | Blaimer et al. (2015) | Conway (1992); Heterick (2017) |
| *Myrmecocystus* | Crop | ~14.1 [19.9 – 10.2] | Van Elst (2021) | Froggatt (1896) Snelling (1976) |
| *Plagiolepis* | Crop | ~11.2 [24 – 3] (low sample size) | Blaimer et al. (2015) | Heterick (2022) |
| *Prenolepis* | Fat body | ~15 [19 – 9] | Boudinot et al. (2022) | Tschinkel (1983) |
| *Proformica* | Crop | ~20 (divergence) | Tinaut and Ruano (2021) | Galkowski (2017) |
| *Zatania* | Crop | ~15 [18.5 – 9] | Boudinot et al. (2022) | Wheeler (1936) |
| *Formica* | Crop | ~17 [21 – 4] | Boudinot et al. (2022) | Cosens & Toussaint (1985) |
| *Leptomyrmex* | Crop | ~43.8 [54 – 35.2] | Barden et al. (2017) |  |
| *Solenopsis* | Fat body | ~39.1 [31.4 – 47.1] | Ward et al. (2015) | Glancey (1973) |
| *Leptothorax* | Crop | [~48 – 25] | Ward et al. (2015) | Børgesen (2000) |
| *Myrmica* | Crop | ~34 | Jansen et al. (2010) | Børgesen (2000) |
| *Monomorium* | Crop | [~50 – 30] | Ward et al. (2015) | Børgesen (2000) |
| *Carebara* | Crop | ~43 [50 – 30] | Ward et al. (2015) | Azorsa & Fisher (2018) |
| *Pheidole* | Fat body | 35.2 [46.6 – 24.9] | Ward et al. (2015) | Tsuji (1990) |

Lineage dates are derived from published molecular-based divergence estimates and indicate the age of the last common ancestor for each genus, except where otherwise noted. Lineage dates denoted with (divergence) correspond with the last common ancestor of the focal genus and its closest living relative sampled in the corresponding phylogeny – such instances reflect inadequate sampling to confidently estimate crown-ages and are therefore overestimates. Instances of single species repletes are excluded; it is not possible to estimate the age of these lineages with current available data. Color legend: Formicinae (orange), Dolichoderinae (green), Myrmicinae (blue).

This study marks the first ancestral state reconstruction of repletes in any genus of ants, and the estimated ages of lineages that contain replete species provide an opportunity to assess the temporal distribution of living food storage (Tab. 1). Molecular-based divergence estimates suggest that crown-group *Carebara* Westwood, 1840, *Leptothorax* Mayr, 1855, and *Monomorium* Mayr, 1855 each originated in the early Eocene, prior to *Leptomyrmex.* Repletism is present but not pervasive in these older taxa, which prevents a clear reconstruction of replete origins – it is not yet known whether repletes evolved once early in the history of these lineages and were subsequently lost in several descendants, or if repletism was recently acquired across multiple distantly related species, for example. Although *Leptomyrmex* is not the oldest lineage to contain repletes, the definitive reconstruction of ancestral repletism here establishes the first clear indication that ants with “honeypot” repletes were present in the Eocene. Future ancestral reconstructions of replete workers across ant lineages will further reveal the tempo of replete evolution as a striking case of morphology-enabled division of labor.

## Acknowledgments

This work was partially funded by an NSF CAREER grant to Barden (#2144915) and was supported by the International Consortium for Honeypot Ant Research, which was partially funded through a Rutgers Global Grant to Khadempour (#FP00024523). Stipend support for Sawh was provided through the NSF HRD-1905142 Bridges to the Doctorate Program.

---

**Supplemental Video 1. Replete workers of *Leptomyrmex relictus.*** Video evidencing mobile replete workers emerging from disturbed nest. Filming took place on the morning of August 20, 2020 (~9:00), at an external temperature of 29.3°C and 27.8°C inside the nest (nest humidity ranging from 30% to 60%).

**Supplemental Video 2. Replete workers of *Leptomyrmex relictus.*** Video evidencing mobile replete workers emerging from disturbed nest. Filming took place on the afternoon of August 20, 2020 (~14:00), external temperature of 34.5°C and internal temperature of 29.4° (nest humidity ranging from 30% to 60%).

